# Impact of climatic changes in the Late Pleistocene on migrations and extinctions of mammals in Europe: four case studies

**DOI:** 10.1101/090878

**Authors:** Mateusz Baca, Adam Nadachowski, Grzegorz Lipecki, Paweł Mackiewicz, Adrian Marciszak, Danijela Popović, Paweł Socha, Krzysztof Stefaniak, Piotr Wojtal

## Abstract

Climate changes that occurred during the Late Pleistocene have profound effects on the distribution of many plant and animal species and influenced the formation of contemporary faunas and floras of Europe. The course and mechanisms of responses of species to the past climate changes are now being intensively studied by the use of direct radiocarbon dating and genetic analyses of fossil remains. Here, we review the advances in understanding these processes by the example of four mammal species: woolly mammoth (Mammuthus primigenius), cave bear (Ursus spelaeus s. l.), saiga antelope (Saiga tatarica) and collared lemmings (Dicrostonyx ssp.). The cases discussed here as well as others show that the migrations, range shifts and local extinctions were the main responses to climate changes and that the dynamics of these climate driven processes were much more profound than it was previously thought. Each species reacted by its individual manner, which depended on its biology and adaptation abilities to the changing environment and climate conditions. The most severe changes in European ecosystems that affected the largest number of species took place around 33–31 ka BP, during the Last Glacial Maximum 22–19 ka BP and the Late Glacial warming 15–13 ka BP.

## INTRODUCTION

The Late Pleistocene was a period marked with multiple climate changes of a magnitude greater than those observed today (Rasmussen et al., 2014). The impact of those fluctuations on Late Pleistocene mammal populations have been intensively studied by examination of changes in spatial distribution of fossil mammalian faunas in time (e.g. Graham et al., 1996; Sommer and Nadachowski, 2006; Lister and Stuart, 2008; Markova and van Kolfschoten, 2008). This period witnessed also a mass extinction on an unprecedented scale not observed in fossil records for millions of years (Sandom et al., 2014; Stuart, 2015). By the end of the Pleistocene, the most of terrestrial megafauna species (heavier than 44 kg) became extinct or their population sizes decreased substantially (Barnosky et al., 2004; Koch and Barnosky, 2006). The course and the timing of these events differ from region to region and the causes of these extinctions are the subject of an on-going debate. Some researchers point only to climate changes, some blame human activities like overhunting and habitat alternation (Alroy, 2001; Sandom et al., 2014) whereas others suggest the combination of these two causes as a main trigger (Barnosky et al., 2004; Prescott et al., 2012; Stuart, 2015).

In recent years two main advances accelerated research in this field. The first one was the increase of accessibility and popularity of the AMS radiocarbon dating. The growing number of directly dated fossils allowed for precise tracking of changes in species' geographic distribution, migrations and dating extinction events (e.g., MacPhee et al., 2002; Stuart, 2005; Sommer et al., 2008, 2014; Pacher and Stuart, 2009; Stuart and Lister, 2012, 2014). It enabled direct correlation between these events and climatic and environmental changes. The second advance was the emergence of the ancient DNA studies. The examination of genetic diversity has added another level of complexity to the research on the Late Pleistocene populations. Ancient DNA facilitated investigations of intraspecific processes, such as population replacements or changes in their diversity; events that usually do not manifest in fossil record. These two methods combined together proved to be the most versatile approach in reconstruction of past population histories (e.g., Campos et al., 2010a, b; Lorenzen et al., 2011; Horn et al., 2014; Palkopoulou et al., 2016).

The main aim of our study is to review the recent progress that was made in the research of four species: woolly mammoth (*Mammuthus primigenius*), cave bear (*Ursus spelaeus* s. l.), saiga antelope (*Saiga tatarica*) and collared lemming (*Dicrostonyx* ssp.) and to discuss its implications for our understanding of the impact of climate changes on migrations and extinctions of mammals in the Late Pleistocene.

## WOOLLY MAMMOTH (*MAMMUTHUS PRIMIGENIUS*)

Elephants (Elephantidae) are the largest terrestrial mammals, with stout body, characteristic long, highly pronounced proboscid or trunk, a combination of nose and upper lip, large ears, column-like limbs, and small tail. The woolly mammoth, represents elephants well adapted to the cold and arid steppe-tundra environment (Maschenko, 2002; Lister et al., 2005). This species was widespread during the Late Pleistocene from Western Europe through the whole northern Asia to the northern part of North America (Kahlke, 2015). The morphological data revealed the woolly mammoths were present in Europe from around 200 ka BP until the end of the Pleistocene (Lister and Sher, 2001; Lister et al., 2005). However, genetic studies indicated that the Late Pleistocene history of *Mammuthus primigenius* was characterized by a complex series of range expansions and contractions, demographic changes and clade replacements (Palkopoulou et al., 2013). Phylogenetic analyses revealed that Holarctic mammoths belonged to three distinct mitochondrial (mtDNA) lineages (Fig. 1). The most widespread lineage I had nearly Holarctic distribution, the lineage II was confined to Central-East Asia while specimens from the lineage III to Europe (Debruyne et al., 2008; Palkopoulou et al., 2013). The divergence of lineages I and II was previously estimated to ca. 1 Ma ago (Debruyne et al., 2008; Gilbert and Drautz, 2008), however, most recent estimations suggest much younger date about 300 ka BP (Palkopoulou et al., 2013). Coalescent simulations suggested that actual split of three mammoth populations took place around 200 ka BP and was followed by a demographic expansion that started around 121 ka BP (Palkopoulou et al., 2013). This expansion coincides broadly with the end of Eemian Interstadial, which suggests that mammoths survived this warm period confined to refugial areas and expanded as climate got cooler at the beginning of Weichselian glaciation (Palkopoulou et al., 2013). Surprisingly, this was not supported by the analyses of the whole paleogenomes, which indicated a much earlier expansion ca. 280 ka years ago and the maximum effective population size during Eemian (Palkopoulou et al., 2015).

**Fig. 1.**
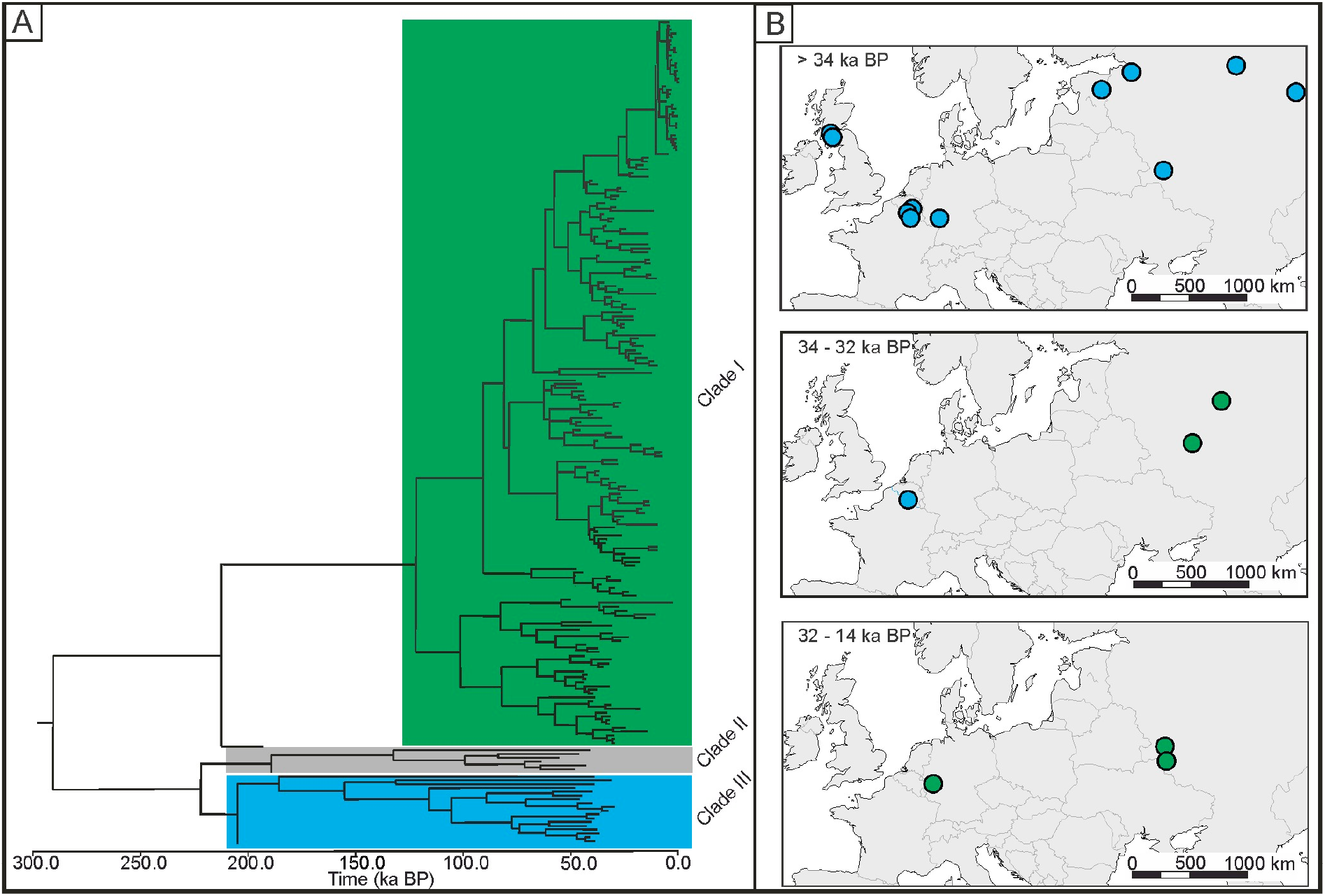
Woolly mammoth (*Mammuthus primigenius*). **A**–Bayesian phylogeny of Holarctic woolly mammoths based on mtDNA cytochrome *b* sequences. The tree is a chronogram where branch lengths denote time elapsed since divergence and the position of tips corresponds to calibrated radiocarbon age of samples; **B**–distribution of paleontological sites with woolly mammoth remains radiocarbon-dated to the indicated periods. Colours indicate mitochondrial DNA lineages (modified after Palkopoulou et al., 2013).

Despite, the ambiguities in the early history of mammoth populations, ancient DNA revealed also two more recent population turnovers. In the Eemian Interglacial and Early Weichselian, woolly mammoths that belonged to clade I were most probably confined to North America. It was estimated that ca. 66 ka BP they started to expand westward through Beringia and reached the area occupied by mammoths from clade II. Both populations lived in sympatry for 20 ka, when the clade II suddenly disappeared from the fossil record ca. 40 ka BP (Palkopoulou et al., 2013). Woolly mammoths belonging to clade I expanded further west and reached Europe. The earliest specimen in Europe originating from this clade came from the site in Vologda Oblast in Russia and was dated to 32 ka BP. The appearance of clade I in Europe coincides with the disappearance of endemic European population (clade III), with the latest specimen dated to ca. 34 ka BP. There is, however, no evidence for any overlap between these populations and it seems that the extinction of clade III was not driven by the appearance of newcomers from the east. This scenario is supported by the lack of radiocarbon dating of mammoth remains in Central Europe between ca. 34 and 32 ka BP, e.g. in Poland (Nadachowski et al., 2011 and Fig. 2). An important increase of mammoth's population size in Europe took place between 31 and 29 ka BP, which is confirmed by dense radiocarbon dating of mammoth remains collected from almost whole North European Plain (Nadachowski et al., 2011; Ukkonen et al., 2011; Markova et al., 2013). Interestingly, during almost the whole Last Glacial Maximum (LGM), between ca. 22 and 18 ka BP (Fig. 2), there are no dated mammoth records in north-western, northern and central Europe sites (Stuart et al., 2004; Nadachowski et al., 2011; Ukkonen et al., 2011), which suggests a long-range contraction. Mammoths re-immigrated to Europe for next 3-4 millennia at the end of the GS-2 (Greenland Stadial 2) with the return of severe climatic conditions and open vegetation (Lister and Stuart, 2008; Nadachowski et al., 2011 and Fig. 2). The next break in the mammoth record coincides roughly with the Bølling-Allerød warming (Greenland Interstadial 1, GI-1) followed by one more recolonization of Latvia and Estonia, dated to the Younger Dryas (GS-1) (Lõugas et al., 2002; Stuart et al., 2002; Ukkonen et al., 2011 and Fig. 7). The last populations of *Mammuthus primigenius* in Europe lived in north-western Russia and disappeared around the Younger Dryas–Holocene boundary ca. 11.8–11.4 ka BP (Lõugas et al., 2002; Ukkonen et al., 2011).

**Fig. 2.**
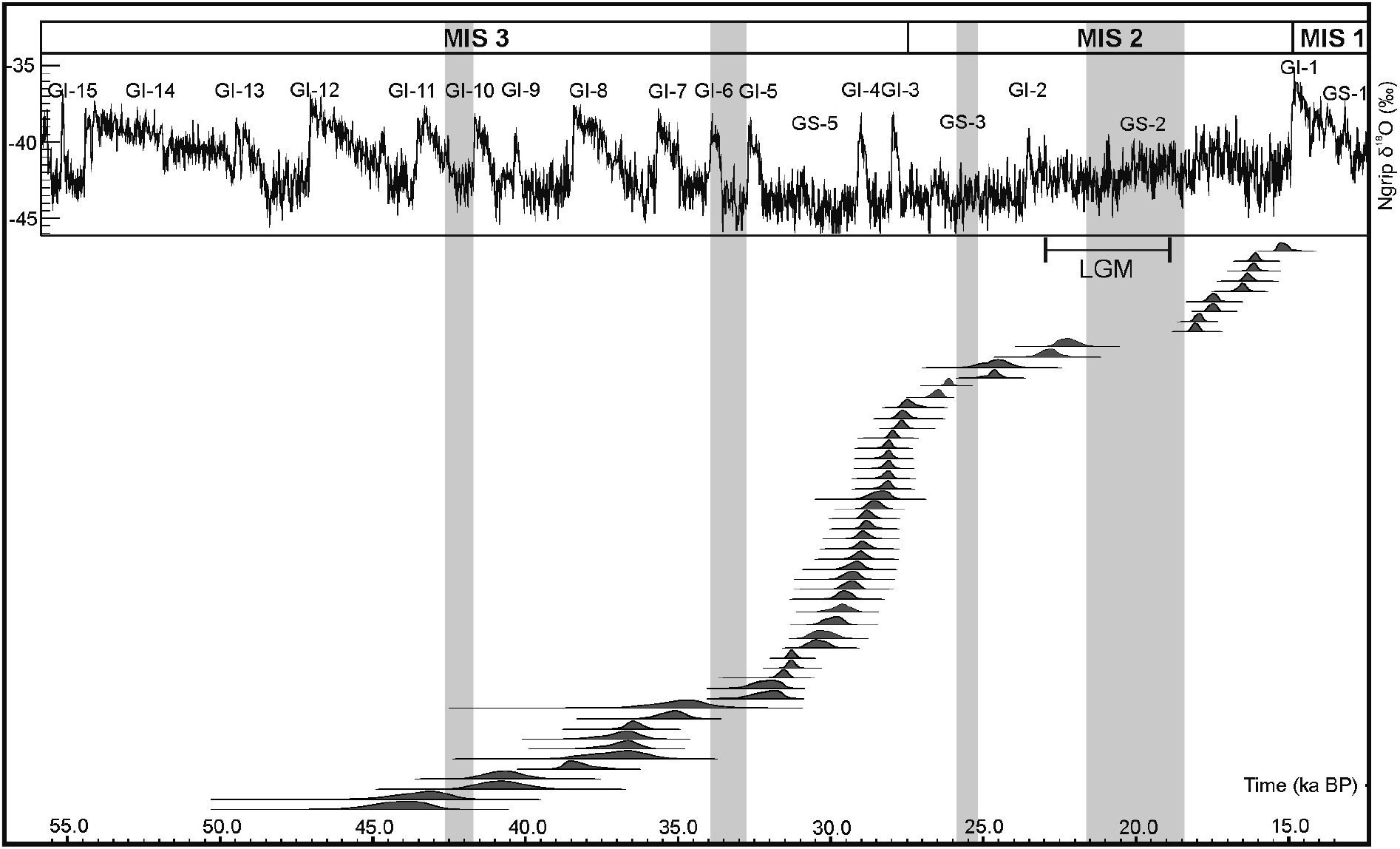
Woolly mammoth (*Mammuthus primigenius*). Radiocarbon dates of woolly mammoth remains from Poland alongside the NGRIP GICC05 ice core δ ^18^O record (Svensson et al., 2008). Dates were compiled from Nadachowski et al. (2011), Pawłowska (2015), Wilczyński et al. (2015), Wojtal and Wilczyński (2015) and recalibrated in OxCal v. 4.2 (Ramsey, 2009) using Intcal13 calibration curve (Reimer et al., 2013). MIS–Marine Isotopic Stages, GS–Greenland Stadials, GI–Greenland Interstadials; LGM–Last Glacial Maximum (after Mix et al., 2001). Grey strips illustrate inferred periods of the absence of mammoths in Poland.

## CAVE BEAR (*URSUS SPELAEUS* S. L.)

The bear family (Ursidae) are large mammals with big head and thick neck, small eyes and short tail, muscular bodies with stout legs and large paws. Most bears are omnivores, although the diets of polar bear (*Ursus maritimus*) and giant panda (*Ailuropoda melanoleuca*) is very narrow and specialized. The former is a carnivore and the latter is an obligate consumer of bamboo. It is assumed that the cave bear (*Ursus spelaeus* s.l.), one of the most widespread mammals in the Late Pleistocene in Europe, was an obligatory vegetarian. It evolved from Middle Pleistocene *Ursus deningeri* and differentiated into several forms recognized at morphological or genetic levels (Hofreiter et al., 2002; Rabeder et al., 2004a, b). In Europe, two main forms, *U. spelaeus* and *U. ingressus* existed. They differentiated probably between 414,000 and 173,000 years ago (Knapp et al., 2009). *U. spelaeus* lived mainly in Western Europe and its remains were found in Spain, France, Germany, Belgium, Italy and Austria, although it was also recorded in Altai (Rabeder et al., 2004b; Knapp et al., 2009). *U. ingressus* inhabited South-eastern and Central Europe and was discovered mainly in Romania, Slovenia, Ukraine, Czech Republic, Poland, Slovakia and Greece but also in Austria, Germany and Switzerland (Rabeder et al., 2004b; Baca et al., 2014). Moreover, two small, dwarf cave bear forms that are considered as a subspecies, *U. spelaeus eremus* and *U. spelaeus ladinicus* were reported in high alpine caves in Austria and Italy (Rabeder and Hofreiter, 2004; Rabeder et al., 2004a) (Fig. 3A). Another major group of large bears, named *U. deningeri kudarensis* was discovered in the Caucasus (Baryshnikov, 1998; Knapp et al., 2009). Recent analysis of mtDNA of the Middle Pleistocene *U. deningeri* from Sima de los Huesos, Atauperca, Spain, revealed that *U. deningeri kudarensis* constitutes the most divergent cave bear lineage (Dabney et al., 2013). It was suggested that it belonged to a separate branch of cave bear evolution and its taxonomic status was changed to *U. kudarensis* (Dabney et al., 2013; Stiller et al., 2014).

**Fig. 3.**
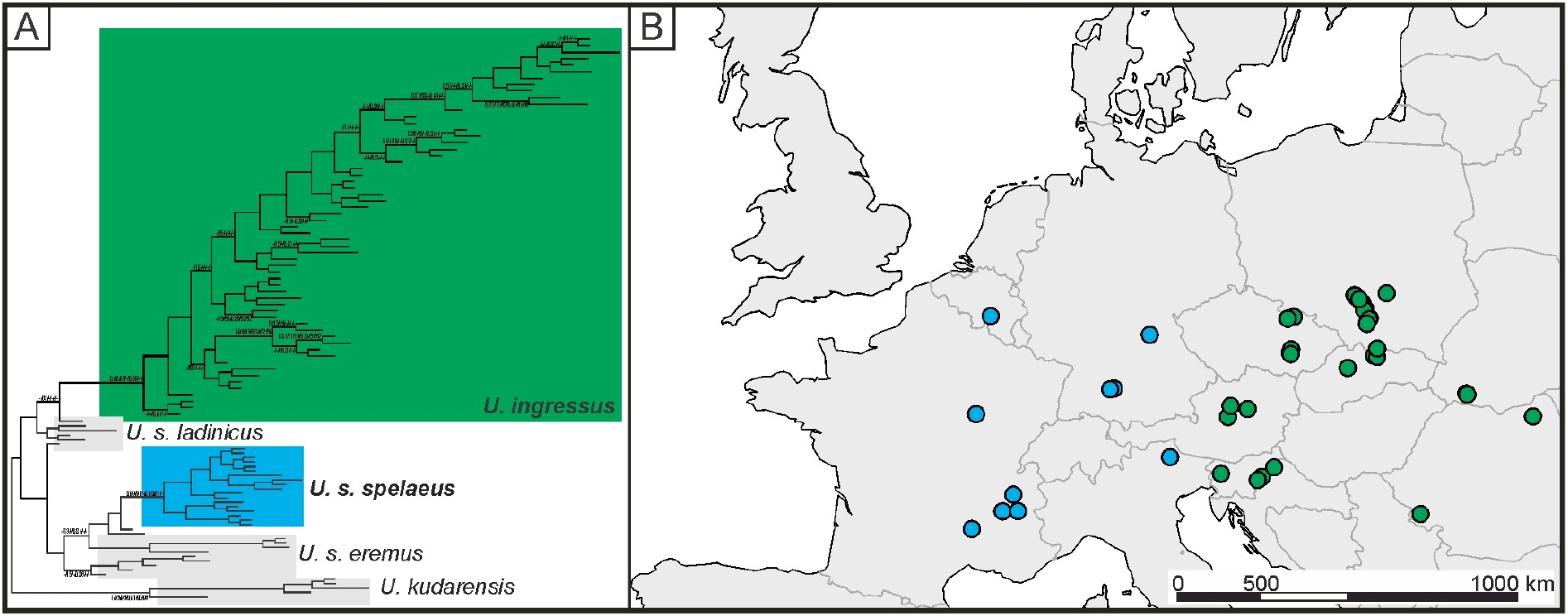
Cave bear (*Ursus spelaeus* s.l.). **A**–Bayesian phylogeny of European cave bears based on mtDNA control region sequences. The tree is a phylogram where branch lengths are proportional to amount of sequence differences; **B**–spatial distribution of cave bear remains classified as *U. s. spelaeus* and *U. ingressus* (after Popović et al., 2015). Colours indicate mitochondrial DNA lineages of cave bears.

The geographic origin of the *U. ingressus* is not known, but the basal position of haplotypes from Romanian site Pestera cu Oase in the phylogeny of this species points to South-eastern Europe (Baca et al., 2012). The phylogeographic picture of *U. ingressus* is unclear as the mtDNA phylogeny lacks significant support and clear phylogeographic lineages are not differentiated (Fig. 3A). It seems, however, that the spread of *U. ingressus* in Europe may have proceeded independently along the main European mountain ridges, Alps and Carpathians (Baca et al., 2014).

It has been proposed that between 60 and 50 ka BP *U. ingressus* started westward migration along the Alps (Hofreiter et al., 2004; Rabeder and Hofreiter, 2004; Münzel et al., 2011). The earliest remains of this species from the Austrian and Swiss Alps, dated to ca. 50 ka BP are known from Gamssulzen cave. The westernmost record of *U. ingressus* came from Schnurenloch site near Bern, Switzerland, but species attribution was based solely on morphology without genetic confirmation (Rabeder and Hofreiter, 2004). Recently, several cave bear specimens from Kraków-Częstochowa Upland, Poland yielded mtDNA haplotypes identical to those found in Alpean *U. ingressus*, which suggests that this population spread also northwards beyond the Carpathian Arc (Popović et al., 2015).

In some cases, the appearance of *U. ingressus* was associated with disappearance of other cave bear forms that inhabited the area. The most interesting evidence of such replacement came from the three cave sites in the Ach Valley in Swabian Jura, Germany: Hohle Fels, Geißenklösterle, and Sirgenstein (Hofreiter et al., 2007; Münzel et al., 2011). The latest occurrence of native inhabitant in this area, *U. spelaeus*, was dated to ca. 31.5 ka BP while the earliest record of *U. ingressus* from Geißenklösterle was dated to 36.3 ka BP. Only a single *U. ingressus* individual yielded such a date while most of others were dated to ca. 32 ka BP, which suggests that the main immigration took place just before the local extinction of *U. spelaeus.* A similar replacement was recorded in Herdengel cave in Austria, where *U. ingressus* replaced one of the small-sized cave bear forms, *U. s. eremus*. All of the *eremus* cave bears were dated by stratigraphic context to more than 60 ka BP, whereas all *U. ingressus* were younger than 37 ka BP (Stiller et al., 2014). The appearance of *U. ingressus* not always resulted in the replacement of other cave bear forms. In the two Austrian caves, Ramesch and Gamssulzen, located about 10 km from each other, *U. ingressus* lived side by side with *U. s. eremus* for at least 15,000 years (Hofreiter et al., 2004).

Less is known about cave bear population that inhabited surroundings of Western Carpathians and Sudetes. Radiocarbon dates suggest more or less continuous presence of cave bears in this area (Nadachowski et al., 2008; Wojtal et al., 2015) during the Late Pleistocene. Preliminary results of genetic investigation of specimens from multiple sites in Poland, Czech Republic, Slovakia and Ukraine revealed that *U. ingressus* was the only form of cave bear present in this part of Europe (Popović et al., 2015; Fig. 3B). It appeared in Carpathians and Sudetes much earlier than in the Alps. The earliest remains confirmed with morphological and ancient DNA analyses are from Niedźwiedzia Cave in Sudetes. Bone collagen from one specimen from this cave was dated with U-Th method to about 80 ka BP (with accompanying ^14^C date >50 ka ^14^C BP) and the other was stratigraphically dated to more than ca. 70 ka BP (Baca et al., 2014). The mtDNA haplotypes of specimens from Niedźwiedzia Cave formed a divergent cluster in phylogenetic trees, which confirms the early separation and expansion of this population (Baca et al., 2012, 2014). Recently, two cave bears with similar mtDNA haplotypes were recorded further west in Zoolithen cave (Upper Franconia, Germany) (Stiller et al., 2014). In this cave, remains of *U. spelaeus* were also discovered and interestingly, an opposite replacement was suggested as the estimated age of both *U. ingressus* specimens was older than of nine *U. spelaeus* individuals (Stiller et al., 2014). These results are, however, based on molecular dating of the remains and should be further confirmed with direct radiocarbon dating.

Another debated issue in cave bear populations' history is the timing and causes of their extinction. Direct radiocarbon dates indicate that last cave bears went extinct prior to the LGM. Until recently, it was thought that they disappeared from fossil record quite synchronously in different parts of Europe around 24 ^14^C ka BP (28 ka BP) at the end of GI-3 (Hofreiter et al., 2002; Pacher and Stuart, 2009; Bocherens et al., 2014). Palaeogenetic analyses showed, however, that the demise of cave bears started ca. 50 ka^14^C BP (Stiller et al., 2010), thus about 25,000 years before their final extinction. It has been proposed that the changing climate was one of the main causes of the cave bear extinction.

Cave bears are generally considered herbivorous based on their craniodental adaptations (Kurtén, 1976; Wiszniowska et al., 2010; van Heteren et al., 2014) and most of the studies of stable isotopes (δ^13^C, δ^15^N) from bone and tooth collagen confirm that they were strict vegetarians (Bocherens et al., 1994, 1997; Taboada et al., 1999; Fernández-Mosquera et al., 2001; Münzel et al., 2011; Krajcarz et al., 2016). The climate change which began since GS-3 caused severe transformations in plant communities all around Europe (Helmens, 2014). Vegetation seasons shortened and the availability of high quality plant material, which seems crucial for cave bears survival, decreased. Dietary habits of cave bears did not change during their presence for the last 10,000 years in Europe and this ecological niche conservatism may have led to decline of their populations (Bocherens et al., 2014).

Beside the environmental changes several other factors might have influenced the cave bear populations. There is a substantial evidence of hunting the cave bears by humans (Münzel et al., 2011; Wojtal et al., 2015), as well as their competition for caves as a shelter (Grayson and Delpech, 2003). Possibly also large carnivores like cave lion (*Panthera spelaea*) and cave hyena (C*rocuta crocuta spelaea*) hunted hibernating cave bears (Diedrich, 2014).

Recently several young cave bear specimens were reported in Western Europe. One specimen from Rochedane site (French Jura), confirmed with ancient DNA as *U. spelaeus,* yielded an AMS date of about 28.5 ka BP (Bocherens et al., 2014). Two specimens from Chiostraccio Cave (Siena, Italy) were dated to ca. 28 and 27 ka BP, respectively (Martini et al., 2014). Stiller et al. (2014) suggested that cave bear populations might have declined from east to west as most of the samples younger than 30 ka BP were found in Western Europe. However, young specimens were also reported in Eastern Europe (Fig. 4B). Two specimens from Kraków-Częstochowa Upland, from Deszczowa and Komarowa caves were dated to 28.6 ka BP and confirmed as *U. ingressus* (Popović et al., 2015; Wojtal et al., 2015). Another one from Isabela Textorisowa cave (Velka Fatra Mts., Slovakia) (Sabol et al., 2014) was similarly dated to 28.7 ka BP. The youngest genetically confirmed cave bear specimen so far come from Stajnia cave in Poland and was dated to around 26.1 ka BP. Baca et al. (2016) gathered 206 radiocarbon dated specimens and following eight approaches estimated the extinction time of *U. spelaeus* sensu lato to between 27.0 and 24.3 ka BP. These findings clearly indicate that the pattern recognised by Stiller et al. (2014) was a result of sampling bias and that the late cave bear survived independently in isolated populations in different parts of Europe even into the middle of GS-3 stadial. Especially the karst regions may have provided suitable microclimate for long survival of this species (Baca et al., 2016).The large amount of directly dated and genetically analysed cave bear remains provided insight into cave bear population dynamics in Europe. However, some aspects of mode of life of cave bears and details about their extinction, evolution and phylogeography still await explanation.

**Fig. 4.**
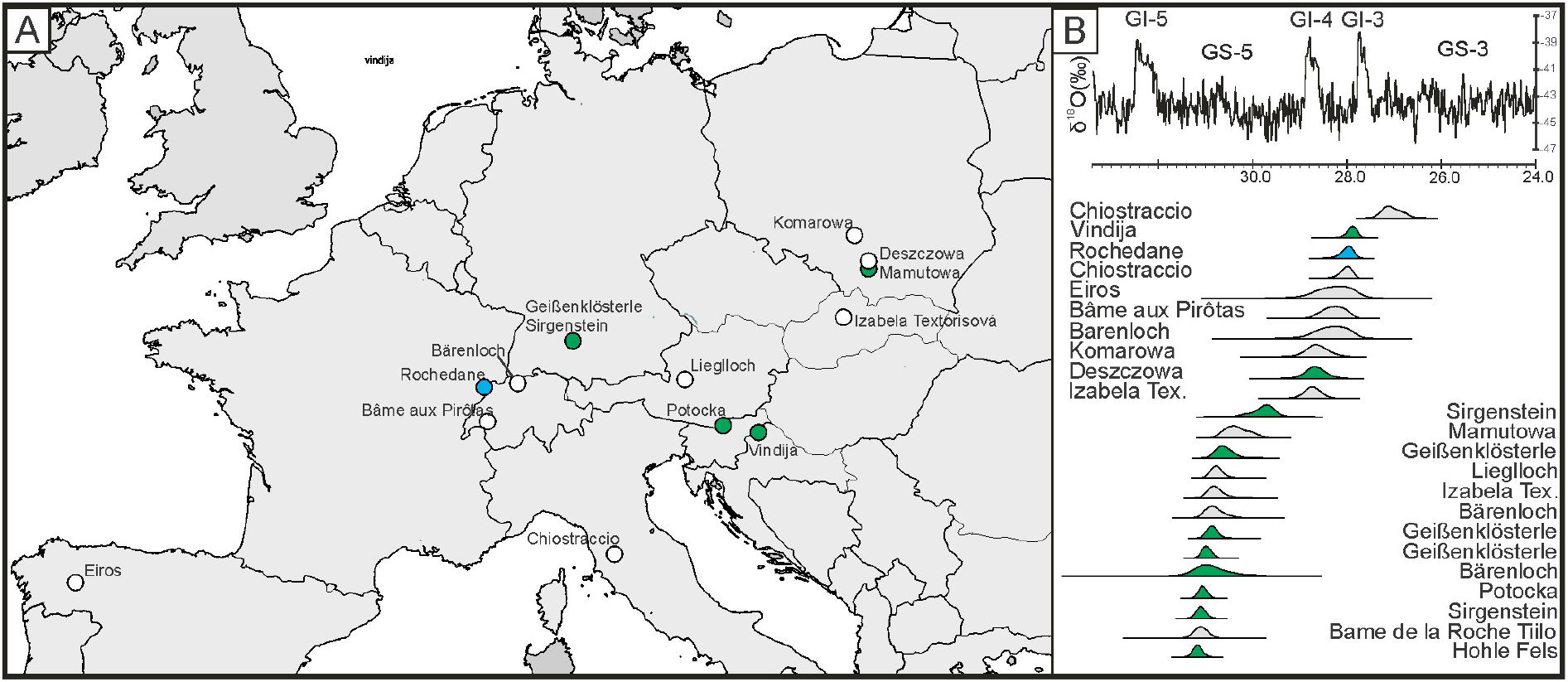
Cave bear (*Ursus spelaeus* s.l.). **A**–distribution of paleontological sites with cave bear remains younger than 32 ka BP, **B**–Cave bear remains younger than 32 ka BP alongside the NGRIP GICC05 δ ^18^O ice core record. Specimens genetically determined as *U. spelaeus* are coloured in blue, whereas *U. ingressus* in green.

## SAIGA ANTELOPE (*SAIGA TATARICA*)

Hollow-horned ruminants (Bovidae) are the most diverse family in the order Artiodactyla. More delicately build species, including the genus *Saiga*, belong to the tribe Antilopini together with gazelles and other antelopes (Groves and Leslie, 2011). The bizarre-looking saigas are not related to sheep and goats, as was used to be thought, but are the sister group of the gazelles. They are non-territorial and nomadic antelopes, gathering in massive herds of many thousands individuals before migrations. Saiga is a specialised steppe herbivore adapted to flat plains and avoiding rugged terrains. At present it inhabits dry steppes and semi-deserts but during the Late Pleistocene it was widespread in vast areas of Eurasia and North America belonging to so-called “*Mammuthus*–*Coelodonta*” faunal complex (Kahlke, 1999, 2014). The number of dispersal events to Europe during the Late Pleistocene was limited by ecological requirements of this species and weather conditions. The high thickness of snow cover in winter was the largest restriction on long-distance migrations of saiga.

There is still hot debate whether saiga represents two or one species. Some palaeontologists distinguish two fossil species: *Saiga borealis* and *S. tatarica* (Baryshnikov and Tikhonov, 1994), others only one species with two subspecies, *S. tatarica borealis* and *S. tatatrica tatarica* in spite of some distinct differences found in the skull morphology between both forms (Kahlke, 1991; Ratajczak et al., 2016). The current genetic studies also suggest that both fossil and recent saiga should be classified as one species, *Saiga tatarica* (Kholodova et al., 2006; Campos et al., 2010b).

The permanent occurrence of this antelope in steppe areas in Eastern Europe north to Black Sea, in the Crimea and Dobruja (Dobrogea) in Romania during the whole MIS 3 cannot be questioned (Markova et al., 1995; Pean et al., 2013; Ridush et al., 2013). Previously published overviews suggest that saiga was also present in Western and Central Europe in MIS 3 (Stewart, 2007; Markova et al., 2010). However, this evidence was not recently confirmed by direct radiocarbon dating (Nadachowski et al., 2016). During MIS 2 and early part of MIS 1 (the Late Glacial), saiga appeared in Europe in three immigration waves (Fig. 5). The oldest one was probably restricted only to Central Europe (e.g. Poland and Czech Republic) (Nadachowski et al., 2016) and was temporally limited to GI-2, i.e. a short warming period between 23.3 and 22.9 ka BP (Rasmussen et al., 2014). The second, geographically much wider migration started just after the LGM, ca. 19.5 ka BP, and lasted for ca. 3,500–4,000 years to ca. 15.5 ka BP (Nadachowski et al., 2016 and Fig. 5). During this time, *Saiga tatarica* was present in the whole Europe north of the Carpathians, Alps and Pyrenees and reached Aquitaine Basin and Gascony in SW France. The comparison of still limited number of direct dates suggests rather migrations of saiga from western Europe to its eastern refugial areas because the dates from France (Langlais et al., 2015) are older than those from Germany and Poland (Nadachowski et al. 2016; Fig. 5). This interesting observation should be, however, confirmed by more direct radiocarbon dating. The reduction of distribution range of saiga in Europe was continued in the Late Glacial (Fig. 5) due to the development of vegetation cover not suitable for this herbivore. However, during cooler episodes of the Late Glacial, between Bølling and Allerød (GI-1d, former Older Dryas) and within the Allerød (GI-1c_2_) (Rasmussen et al., 2014) *Saiga tatarica* probably returned to Central Europe, but this reimmigration was restricted only to the area north of the Carpathians (Nadachowski et al., 2016 and Fig. 5). It seems that this species was unable to extend its range to the west of Europe during the Younger Dryas (GS-1), the last cold phase of the Pleistocene, even the climatic and environmental conditions were suitable for this herbivore.

**Fig. 5.**
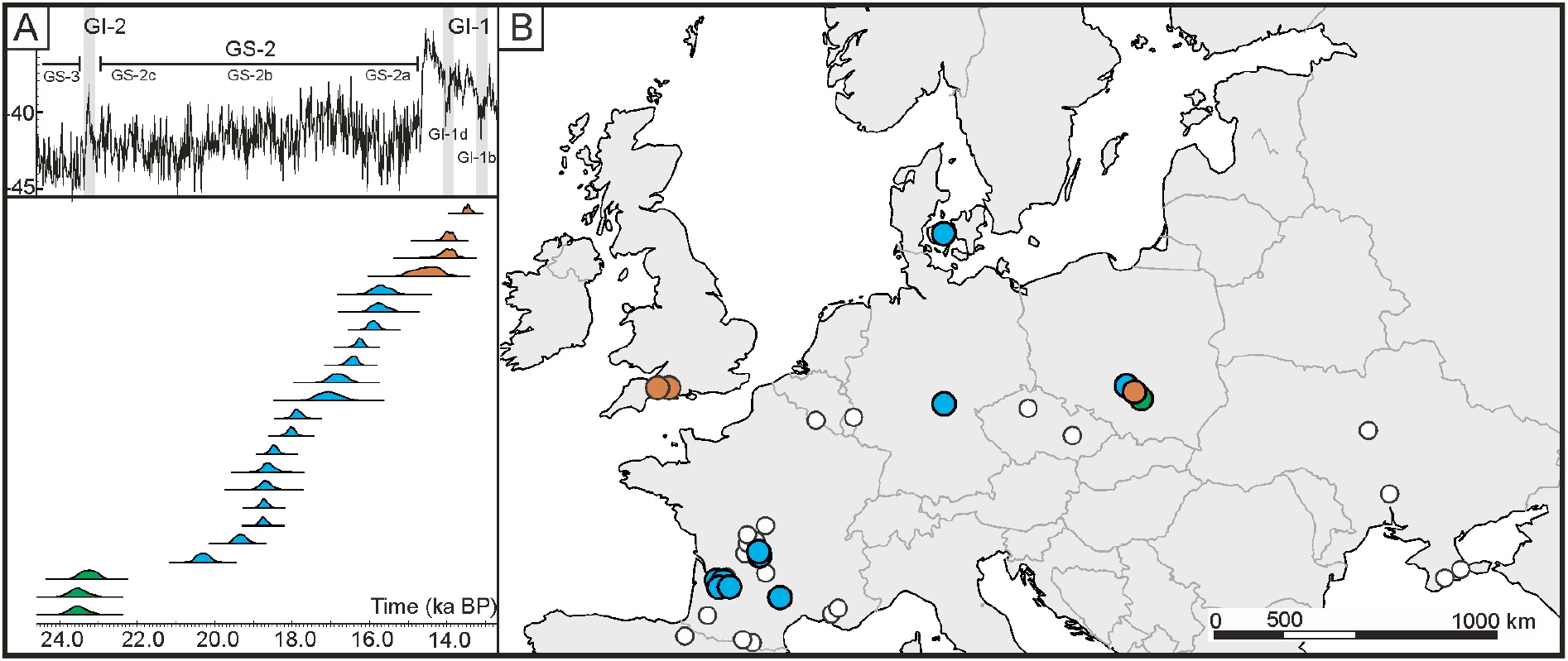
Saiga antelope (*Saiga tatarica*). **A**–a plot of direct radiocarbon dates of saiga antelope in Europe (Langlais et al., 2015 and Nadachowski et al., 2016) alongside the NGRIP GICC05 δ ^18^O ice core record, calibrated using program OxCal v. 4.2 (Ramsey, 2009). GS–Greenland Stadials, GI–Greenland Interstadials. **B**–spatial distribution of sites with saiga remains in Europe. White circles denote sites with remains dated by archaeological context, while coloured circles indicate sites with radiocarbon dated remains. Colours correspond to three putative waves of migration of saiga into Europe (supplemented and modified after Nadachowski et al., 2016).

The history of saiga in the European Late Pleistocene, especially directions of migrations (westward or eastward), needs further investigation basing on new radiocarbon dating and palaeogenetic studies to explain the complex migration events of this endangered ruminant.

## COLLARED LEMMING (*DICROSTONYX* SSP.)

Morphological and molecular evidence shows that voles and lemmings create a monophyletic group of rodents ranked by most authors as Arvicolinae subfamily (Chaline et al., 1999). Collared lemmings (*Dicrostonyx* ssp.) are cold-adapted animals that are restricted to dry and treeless Arctic tundra environments (Kowalski, 1995). They are a key species in trophic networks in Arctic ecosystems as they are prey for predators such as arctic fox, snow owl or stoat. Four contemporary species are recognised within the genus, *D. torquatus* that inhabits Eurasia from White Sea to Bering Strait, as well as *D. groenlandicus, D. richardsoni* and *D. hudsonius* that inhabit North America. These species have been recognised on the basis of phylogenetic analysis of mitochondrial DNA, karyotype diversity and hybridisation experiments (Jarrell and Fregda, 1993; Fedorov and Goropashnaya, 1999). Such classification is currently the most commonly accepted.

At present collared lemmings show nearly circumpolar distribution and do not exceed latitude 65° N, but in the Pleistocene their range was much wider and their remains are known from paleontological sites throughout Europe and Asia (e.g. Markova et al., 2010; Ponomarev and Puzachenko, 2015). Pleistocene collared lemmings are classified according to the growing complexity of occlusal surface of molar teeth. The oldest ones are known from the Early Pleistocene and are classified as *Predicrostonyx* (Nadachowski, 1992), which was followed by *D. renidens, D. meridionalis* and *D. simplicior* in the Middle Pleistocene (Zazhigin, 1980; Smirnov et al., 1986) and *D. gulielmi* in the Late Pleistocene. It was suggested that the transition between *D. gulielmi* and the recent *D. torquatus* took place in the Late Glacial about 15–13 ka BP. Morphometric analyses of molar teeth revealed that during this time, a substantial change in the structure of collared lemming populations took place (Smirnov, 2002). It was further supported by the genetic analyses of contemporary collared lemming populations. Phylogeographic analyses revealed the existence of five allopatric populations within the present distribution of the species (Fedorov et al., 1999). Each of the populations was characterised by low genetic diversity, which was interpreted as an effect of the regional population reductions that might have taken place during the warm periods in Holocene. Another evidence came from the palaeogenetic analyses of collared lemmings remains from Pymva Shor site in northern Pre-Urals that revealed the signature of severe population bottleneck around 14.5 ka BP (Prost et al., 2010). The climate warming during the Bølling/Allerød (GI-1a-c) period in the Late Glacial was accompanied with northward expansion of forests and supposedly this forced the contraction and isolation of collared lemming populations that resulted in their decline (Fedorov, 1999).

Recently, Palakopoulou et al. (2016) used ancient DNA and radiocarbon dating to reconstruct the Late Pleistocene evolutionary history of collared lemmings. They obtained mtDNA sequences from more than 300 collared lemming specimens from multiple palaeontological sites, and 48 direct radiocarbon AMS dates. Phylogenetic analyses revealed that five distinct populations of collared lemmings, represented by five mitochondrial lineages (EA1–EA5, Fig. 6A), existed in Europe. Each of the populations was widely distributed ranging from Western France to Ural Mountains. The radiocarbon dating revealed that subsequent populations succeeded each other through time during the last ca. 50 ka. This was interpreted as a series of collared lemming population extinctions and recolonizations that took place across the whole European continent. The found phylogenetic pattern indicated that subsequent recolonizations proceeded in east to west direction from a hypothetical refugial area in north-eastern Siberia. Dates for two earliest populations (EA1 & EA2) are close to radiocarbon dating limits but it seems that both were already present in Europe prior to 50 ka BP. The EA1 population vanished around 50 ka BP and EA2 population around 42.3 ka BP. Population EA3 appeared in Europe ca. 32 ka BP and disappeared 22.8 ka BP and was followed by population EA4, dated to a short period between 22.2 and 20.5 ka BP. Population EA5 emerged ca. 20.3 ka BP and disappeared from Europe ca. 14.5 ka BP. All Holocene and modern collared lemmings from Siberia belong to this EA5 mtDNA lineage (Fig. 6B).

**Fig. 6.**
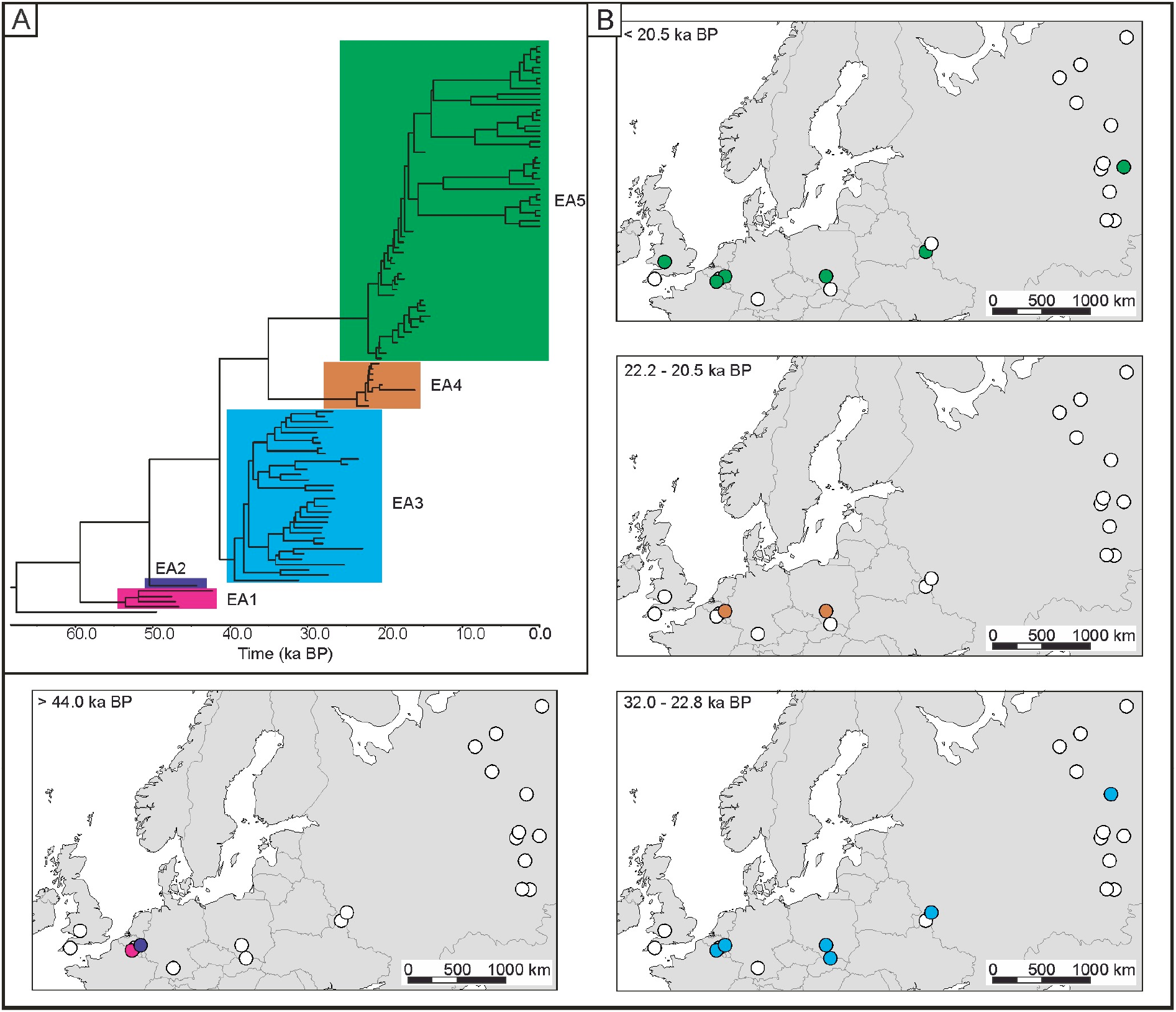
Collared lemming (*Dicrostonyx* ssp.). **A**–Bayesian phylogeny of Eurasian collared lemmings based on mtDNA cytochrome *b* sequences. The tree is a chronogram where branch lengths denote time elapsed since divergence and the position of tips corresponds to calibrated radiocarbon age of the sample. Colours indicate different mtDNA lineages; **B**–Spatial distribution of paleontological sites with collared lemming in different periods. Colours indicate different mtDNA lineages (modified after Palkopoulou et al., 2016).

It was postulated that population turnovers that took place across such a large geographical range, has to be driven by environmental or ecological changes in the steppe-tundra ecosystem. The dating of two earliest populations (EA1 and EA2) was not very precise and the connection between their demise and climate changes remains unclear. The spread of population EA3, which seems reliably dated, started just after GI-5 and might be associated with climate cooling that begun with the onset of the GS-5. Surprisingly, the two following population turnovers, i.e. the replacement of population EA3 by EA4 and EA4 by EA5, took place within the LGM. During this period, arctic tundra environments preferred by the collared lemmings, spread over the large areas of Eurasia (Tarasov et al., 2000). Thus, the causes of these replacements remain unclear, however, it was shown that they coincide with short term warming periods recorded in high resolution palynological record from Lago Grande di Montichio, Itally (Palkopoulou et al., 2016).

## DISCUSSION AND CONCLUSIONS

The high resolution reconstructions of migrations and extinctions events during the Late Pleistocene obtained in recent years showed that species responded to climate changes according to their individual adaptations and there are no grounds for considering species as faunal complexes responding simultaneously and in the same way (Hofreiter and Stewart, 2009; Stewart et al., 2010). This is well illustrated in the comparison of three species reviewed here: woolly mammoth, saiga antelope and collared lemming. They all are considered as members of “*Mammuthus*–*Coleodonta*” faunal complex (Kahlke, 1999, 2014), but their Late Pleistocene histories in Europe differ substantially. During the last 50 ka mammoths were present in Europe more or less continuously, with possible range contraction only during the LGM. The first confirmed appearance of saiga antelope was most probably related with the short term warming GI-2 dated to ca. 23 ka BP. However, only after the LGM, important extension of its range in Europe was observed. Collared lemmings most probably occurred, as mammoths, continuously since 50 ka but their population turnover was much more intensive and happened often than in the case of previous species. Interestingly, along with these apparent differences, some similarities are visible. It seems that most of the observed events are grouped around three periods. The first one covers the GI-5 warming and the onset of the GS-5 cooling around 33–31 ka BP, the second one partly overlaps the maximum extent of Scandinavian Ice Sheet during the LGM ca. 22–19 ka BP and the last one is the abrupt climate warming during the Late Glacial ca. 15–13 ka BP (Fig. 7). In agreement with that, the climate during the LGM (as defined by Mix et al., 2001) and the warming during the Late Glacial have been previously recognized as main factors that substantially influenced the distribution of many mammal species in Europe.

**Fig. 7.**
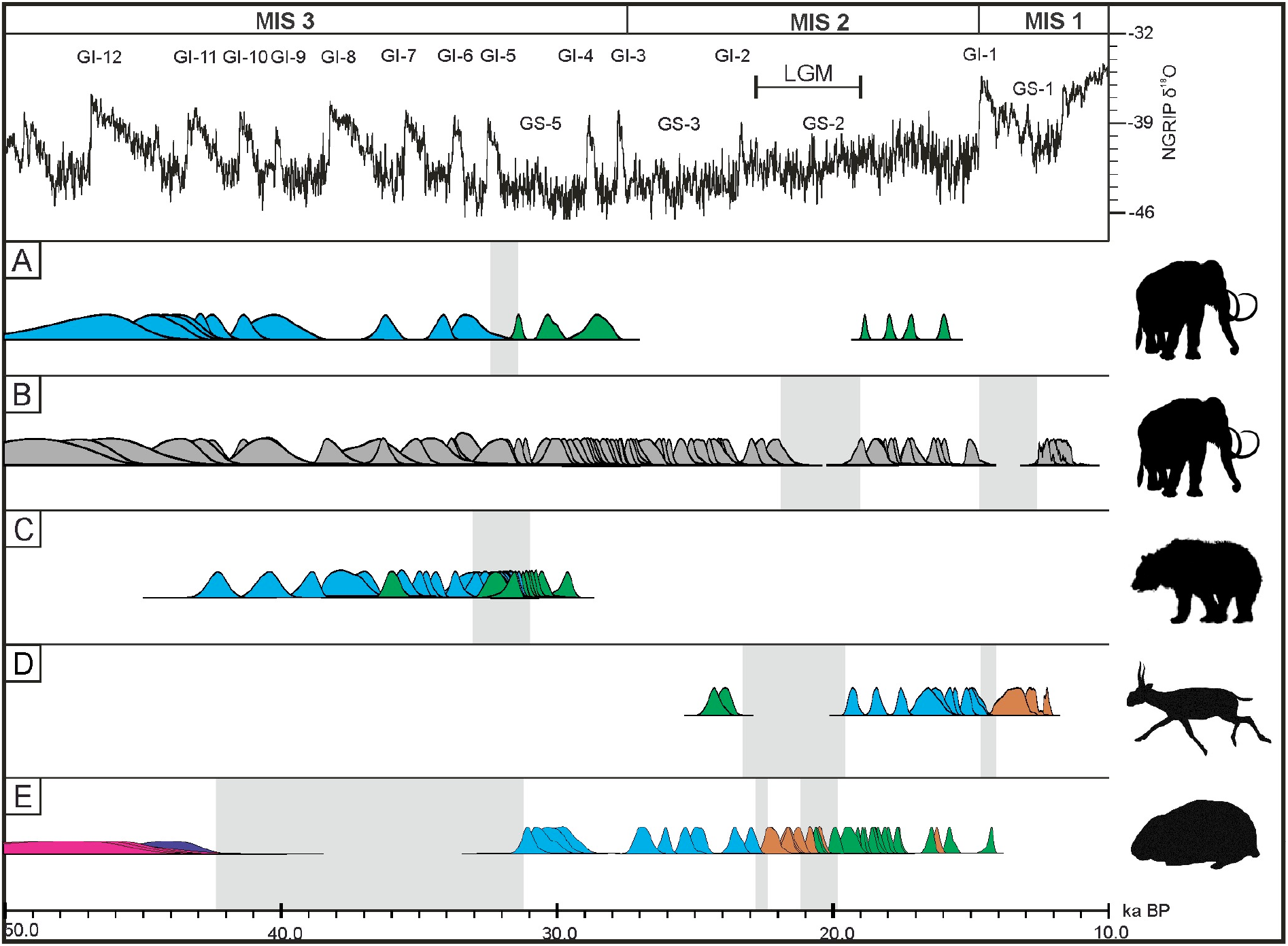
Population turnovers and extinctions in Europe based on radiocarbon dating. **A**–woolly mammoth, colours indicate different mtDNA clades (only specimens with genetic information are presented); **B**–woolly mammoth from Europe, without the differentiation into the clades, source of the dates is like in Fig. 2 supplemented with data from Nadachowski et al. (2011) and Ukkonen et al. (2011); **C**–cave bear, blue–*U. s. spelaeus*, green–*U. ingressus*; **D**–saiga antelope; **E**–collared lemmings, colours indicate different populations (EA1–EA5). MIS–Marine Isotope Stages, GI–Greenland Interstadials; GS–Greenland Stadials; LGM–Last Glacial Maximum (after Mix et al., 2001). Vertical strips indicate timing of population turnovers or extinctions.

During the LGM, numerous species withdrew from Europe or restricted their ranges into so-called southern glacial refugia (Hewitt, 2004; Stewart and Cooper, 2008). This concerns most of temperate species like red deer (*Cervus elaphus*) (Sommer et al., 2008; Skog et al., 2009), roe deer (*Capreolus capreolus*) (Sommer et al., 2009; Sommer and Zachos, 2009), brown hare (*Lepus europaeus*) (Stamatis et al., 2009) as well as cold adapted ones, like mammoths and saiga antelopes (Fig. 7). During the maximum extent of Scandinavian Ice Sheet, environmental conditions in Central and North-western Europe were extremely harsh, with continuous permafrost covering most of Poland, Germany, Belgium, Netherlands and northern France (Huijzer and Vandenberghe, 1998). Ukkonen et al. (2011) suggested that during that time productivity of environments was extremely low, which forced the contraction of even cold adapted species. However, the harsh conditions during the LGM do not seem to lead to many extinction events. Cooper et al. (2015) pointed that, the gradual climate deterioration prior to and the relative stability during the LGM allowed mammalian populations to retreat or to adapt to the changing environment. Two Europe-wide replacements of *Dicrostonyx* populations observed between 23 and 20 ka BP suggest, however, that some short-term environmental changes must have taken place at that time.

The Late Glacial period was marked with several climate oscillations, with warm Bølling-Allerød Interstadial (GI-1) interrupted with Older Dryas (GI-1d) and followed with Younger Dryas (GS-1) cold phases (Steffensen et al., 2008). These rapid changes are linked with the extinction of cave lions (*Panthera spelaea*) (Stuart and Lister, 2011) and woolly rhinoceros (*Coelodonta antiquitatis*) (Stuart and Lister, 2012).

The period of 33-31 ka BP(33-1 ka BP) has been put forward only recently with the analyses of collared lemmings. Palkopoulou et al. (2016) noticed that the appearance of one of the *Dicrostonyx* populations in Europe on the onset of GS-5, ca. 31 ka BP, coincides broadly with the replacement of the endemic European mammoth population by another one from Asia and with the replacement of *U. s. spelaeus* by *U. ingressus* in Central Europe (Fig. 7.). Furthermore, the recent new radiocarbon AMS data showed that the last populations of spotted hyena (*Crocuta crocuta*) also became extinct ca. 31 ka BP (Stuart and Lister, 2014). The recent worldwide survey of extinction and population replacements revealed that timing of these events was significantly correlated with rapid and high amplitude climate changes during interstadials (Cooper et al., 2015). The concentration of such events around the GI-5 fits well to this hypothesis as this interstadial represents one of the most instant climate changes during last glaciation.

All four case studies presented here unveil the Late Pleistocene communities as complex and highly dynamic, characterized by geographic range shifts, migrations, replacements and local extinctions, which seem to be common phenomena rather than exceptions. Cooper et al. (2015) note that such processes might be essential for maintaining ecosystems stability in the periods with high climate variability.

## Acknowledgements

The work related to cave bear was supported by Polish National Science Centre grant no. 2012/07/B/NZ8/02845. We are grateful to the reviewers Lembi Lõugas and Thijs van Kolfschoten for their constructive comments and to Martyna Molak for her suggestions and help with proofreading and correction to the manuscript.

